# Single Cell atlas of uterine myometrium and leiomyomas reveals diverse and novel cell types of non-monoclonal origin

**DOI:** 10.1101/2020.12.21.402313

**Authors:** Jyoti Goad, Joshua Rudolph, Jian-Jun Wei, Serdar E Bulun, Debabrata Chakravarti, Aleksandar Rajkovic

## Abstract

Uterine leiomyomas are the most common tumors of the female reproductive tract with significant morbidity that includes excessive bleeding, infertility and pregnancy complications. The origin and cellular composition of leiomyomas is controversial, yet very important in better understanding the pathogenesis of these tumors. We applied single-cell RNA sequencing to better understand cellular heterogeneity of uterine leiomyomas and normal myometrium at the molecular level. Our data reveal previously unknown heterogeneity in the smooth muscle cells, fibroblast cells, and endothelial cells of normal myometrium and leiomyomas. We discovered a novel lymphatic endothelial cell population in uterine leiomyomas and that the immune as well as transcriptional profile of leiomyomas is *MED12* genotype-dependent. Moreover, we show that leiomyoma cell moiety is not monoclonal in nature. Our work describes unprecedented single cell resolution of normal uterine myometrium and leiomyoma tumors and provides insight into tumor specific hormone responsiveness and extracellular matrix accumulation.

## Introduction

Uterine leiomyomas, also known as fibroids, are benign tumors of the myometrium affecting nearly 25% of women in their reproductive age^1^. These tumors significantly affect women’s quality of life and are the single most common cause of hysterectomy^1^. Each year, approximately 300,000 myomectomies and 200,000 hysterectomies are performed in the United States to remove either leiomyoma tumors or the whole uterus^2,3^. Despite its importance to women’s health, there are currently no leiomyoma specific therapeutics. Moreover, the underlying origin and heterogeneity of leiomyomas continues to evolve.

Extensive genetic studies from our group and others have shown *MED12* exon 2 as a hotspot for genetic variants that associate with leiomyoma in about 70% of the cases^4–6^. In contrast, the *MED12* variant negative leiomyomas have a highly heterogenous genomic landscape^7,8^. Studies have shown that *MED12* variant negative leiomyomas are bigger than leiomyomas expressing *MED12* variant allele^9^. The genotype dependent difference in leiomyoma size is due to differences in the cell composition, rate of proliferation, and accumulation of extracellular matrix^10^. Leiomyomas are considered monoclonal tumors of smooth muscle cell in uterus^11^. However, recent histological and flow cytometry studies have shown the presence of smooth muscle cells, fibroblasts, endothelial cells and immune cells in leiomyomas^12^. How these cells contribute to the leiomyoma formation and the molecular salient characteristics of the cellular heterogeneity in leiomyomas remain undetermined.

Here we utilized single-cell RNA sequencing to understand the underlying cellular heterogeneity in normal myometrium and uterine leiomyomas. We have identified two novel lymphatic endothelial cell populations that are present in uterine leiomyomas. We also determined the transcriptomic changes in the leiomyoma cell clusters based on the genetic mutation. Our data shows that different smooth muscle and fibroblast cell clusters expand in leiomyomas as compared to myometrium. Immune cell infiltration differs with the genotype of the leiomyomas. Our work has identified unanticipated cellular differences in leiomyomas and the myometrium, which could help the future direction of developing targeted therapy for leiomyoma treatment.

## Methods

### Study subjects

To resolve the cellular identity of the cells, we collected 66,339 cells from the leiomyomas and the myometrium from a total of 8 patients. Five out of the 8 samples were matched which were collected from patients undergoing hysterectomy for the treatment of leiomyomas. The non-matched leiomyomas were collected from patients undergoing myomectomy. Histopathological assessment of the collected samples confirmed the identity of the samples.

### Patient tissue collection and genotyping

Uterine leiomyomas and normal myometrium were collected from the patients undergoing hysterectomy or myomectomy with informed consent. The study was approved by the UCSF institutional review board, ethics approval 17-22669. Fresh tissue samples were collected immediately after the surgery, placed in ice cold DMEM/F12, and immediately sent to the lab. A small part of each tissue obtained was snap frozen to perform DNA isolation for genotyping as described previously by us^4^. Additionally, a part of the sample was fixed in formalin overnight for histology. Patient information is provided in S table 1.

### Preparation of single cell suspension from the fresh tissues

The tissue samples were collected and washed in HBSS (Sigma). Leiomyomas and myometrium were cut into pieces of 3-4 mm. These pieces were then added to 3-4 ml of digestion media containing 0.1 mg/ml Liberase (Roche, 501003280), 100 U/ml DNase I (Sigma, D4527), and 25 U/ml Dispase (Sigma, D4818) in DMEM (Life Technologies, 12634010) per gram of the tissue and mechanically dissociated using gentle MACS dissociator (Miltenyi Biotech, Germany) for 30 mins at 37°C to prepare a single cell suspension. The cell suspension was then pipetted up and down with 25 ml,10 ml, and 5 ml pipettes for 1 minute each and then filtered through 70 μm filter (Corning, 431751). Debris was then removed from the cell suspension using debris removal solution (Miltenyi Biotech, 130-109-398) as per the manufacturer’s instructions. The cells were then incubated with RBC lysis buffer (Thermofisher Scientific, 00-4333-57) for 5 mins on ice to remove the red blood cells. The cells were then resuspended in PBS containing 0.4% ultrapure BSA (Thermofisher Scientific, AM2616) and passed through the 70-μm cell strainer (Bel-Art, H13680-0070) to obtain single cell suspension.

### Immunofluorescence and RNAscope

Immunofluorescence was performed as described previously^13^. Briefly, tissue sections were incubated with the following primary antibodies: CD8a (1:200, Cell signaling), CD20 (1:250, Abcam), *NUSAP1* (1:1000, Abcam), *PDPN* (1:200, Cell signaling), overnight at 4°C, then incubated with secondary antibody for 1 hour at room temperature. For immunofluorescence, the images for all of the samples, were taken at the same exposure using a Nikon microscope (Nikon, Japan).

RNAScope Fluorescent multiplex assay was performed using the RNAScope Multiplex Fluorescent v2 kit (Advanced Cell Diagnostics) as per the manufacturer’s instructions. Images were obtained using the Leica SP8 confocal microscope (Leica Biosystems, Germany).

### Dual immunofluorescence and in-situ hybridization

Dual immunofluorescence and in-situ hybridization were performed using the RNAScope Multiplex Fluorescent Reagent kit v2 (Advanced Cell Diagnostics) as per the manufacturer’s protocol. Briefly, after the RNAScope protocol, the tissue sections were blocked overnight at 4°C. Tissues were then incubated with α-sma (1:200, Sigma Aldrich) antibody for an hour at room temperature followed by incubation with the secondary antibody for 1hr at room temperature. Images were obtained using the Leica SP8 confocal microscope (Leica Biosystems, Germany).

### Single cell sequencing

Single cells were processed through the 10X Chromium system (10X Genomics, USA) using single cell 5’ library and Gel bead kit (10X Genomics) as per the manufacturer’s instructions. Briefly, the single cell suspensions were partitioned into gel bead-in-emulsions which were utilized to generate the barcoded cDNA libraries. The single-cell barcoded cDNA libraries were then sequenced using an Illumina NovaSeq 6000 sequencing system (Illumina, USA).

### Pre-processing scRNA-seq data

Cellranger v.2.1.0 single-cell software suite from 10X Genomics was used to demultiplex fastq files, align reads to the Genome Reference Consortium Human Build 38 (hg38) transcriptome, and extract cell and UMI barcodes. Raw cell count by transcript matrices were imported into R and Seurat for analysis^14^. Each sample’s expression matrix was filtered to remove low-quality cells, defined as having fewer than 200 reads, greater than 2500 reads, or more than seven percent mitochondrial gene expression. Samples were merged using Seurat’s integration anchors. Any cells with more than one percent hemoglobin gene expression were removed, with hemoglobin genes considered as *HBA2, HBA1*, and *HBB*. The remaining hemoglobin gene expression was regressed out from the expression matrix using Seurat’s scale.data feature. To counteract differences in sequencing depth across cells, transcript counts were normalized in each cell to transcripts per 10,000 unique molecular identifiers (UMI).

### Dimensionality reduction, clustering, and differential expression analysis

Seurat was used to cluster the merged object into subsets of cells. This workflow includes finding variable genes, running principal component analysis on variable genes, running Uniform Manifold Approximation and Projection on Principal components (UMAP) (1:20), and graphed clustering using KNN and Louvain clustering (with Seurat FindClusters () resolution 0.4). Cell types were defined empirically using expression of marker genes. Cells were partitioned into smooth muscle, fibroblasts, endothelial, and immune cells based on the known signature markers. The following signature markers were used to identify the cell clusters: smooth muscle cells (*MYH11, TAGLN, ACTA2, CNN1, DES, CALD1*), fibroblast cells (*VIM, ALDH1, CD90, FN1, DCN, OGN, MGP, COL1A1, COL1A2, COL3A1*), endothelial cells (*PECAM1, CD31, CDH11, VWF*), and immune cells (*CD3D*, *CD3E, FCER1G, MS4A1, CD79B, CST7, GZMB, FCGF3A, MS4A7*).

Further clustering was performed for each cell type based on the expression of the signature markers (with resolution 0.4) to delineate heterogeneity. For all clusters heatmaps, *t-* distributed stochastic neighbor embedding (*t*-SNE) and UMAP visualizations, violin plots and dot plots were produced using Seurat functions with ggplot2, pheatmap and grid R packages. Differential gene expression analysis was performed using Seurat based on the non-parametric Wilcoxon rank sum test.

### Batch effect and quality control

Cell barcode and gene matrices were constructed using Cellranger and analyzed in R with Seurat^14^. Cells with fewer than 200 features and genes in less than 3 cells were filtered. To account for potential doublets and low-quality cells in the analysis, only cells with fewer than 2500 features and less than 7 percent total mitochondrial genes were used for downstream analyses. Cell samples were then combined using the Seurat standard workflow to integrate cells. *HBA2, HBA1*, and *HBB* genes were used to calculated hemoglobin gene percentage, some cells with high percent were removed, and the remaining effect was regressed out using the Scale.Data feature. Erythrocytes were also removed from downstream analysis.

### Standard Error Barplots

Percentages of samples were used instead of raw cell numbers to normalize for the varying number of cells introduced to the experiment by each sample. For each subset of cells, cell proportions were calculated as a percentage of the number of cells in a cluster from a sample, divided by total number of cells from the same sample. Standard error of the mean was then calculated for each cluster of samples and median percentages per cluster were used to plot.

### Volcano Plots

Seurat’s FindMarkers with default parameters was used to get the fold change between *MED12* positive leiomyomas and myometrium as well as *MED12* negative leiomyomas and myometrium. All data was compiled and plotted in R with ggplot.

### Clonality analysis

Dsc-pileup in the freemuxlet tool was used to construct barcode and variants table^15,16^. Variants from cells which weren’t filtered by preprocessing were used in conjunction with Seurat meta data to print UMAPs with variant information.

### Statistical Analysis

Dirichlet multinomial regression was utilized to determine the statistical confidence that proportions of conditions in each cluster are not a product of random sampling. A Fisher t-test was also leveraged to compare cell numbers from one condition to another in each cluster. Bar plots show means of sample percentages, with standard error of the mean confidence intervals.

## RESULTS

### Single cell atlas of the human myometrium and leiomyomas

We collected a total of 66,339 cells present in normal myometrium and leiomyomas from a total of eight patients (Fig. 1A). After accounting for technical and biological variation, clustering of the 34,435 high quality cells (11,235 from myometrium (n=5), 15,417 from *MED12* variant positive (n=5) and, 7,783 cells from *MED12* variant negative leiomyomas (n=3)) revealed the presence of 18 different clusters across known cell lineages (Fig. 1B, C). These cell clusters were highly reproducible as all samples were represented in nearly all clusters (Fig. 1D). The subpopulations included known cell types previously identified through histology and flow cytometry^12^. These included: smooth muscle cells, fibroblasts, NK cell, T cells, B cells, myeloid cells, and endothelial cells, (Fig. 1C). The annotations were performed using the differential gene expression analysis supported by known gene markers such as *ACTA2, CNN1* for smooth muscle cells (SMC), *VWF* and *PECAM* for endothelial cells (Endo), *PDPN* lymphatic endothelial cells*, DCN* and *LUM* for fibroblasts, *CD3D* for T cells, *MS4A1* and *CD79A* for B cells, *GNLY* and *NKG7* for NK cells, *CD14*, and *S100A8* for myeloid cells, (Fig. 1E, S Fig. 1B,C). All cell clusters were present in most patient samples across all three types of tissue samples: myometrium, *MED12* variant positive leiomyomas and *MED12* variant negative leiomyomas (Fig. 1D, S Fig. 1A). The contaminating endometrial cells (*EPCAM+* or *KRT8+*) and red blood cells expressing HBB-A/ HBA-2 were removed from the datasets from further analysis. Quality control matrices were highly reproducible across patient samples and conditions (S Fig. 1D).

**Fig.1:**
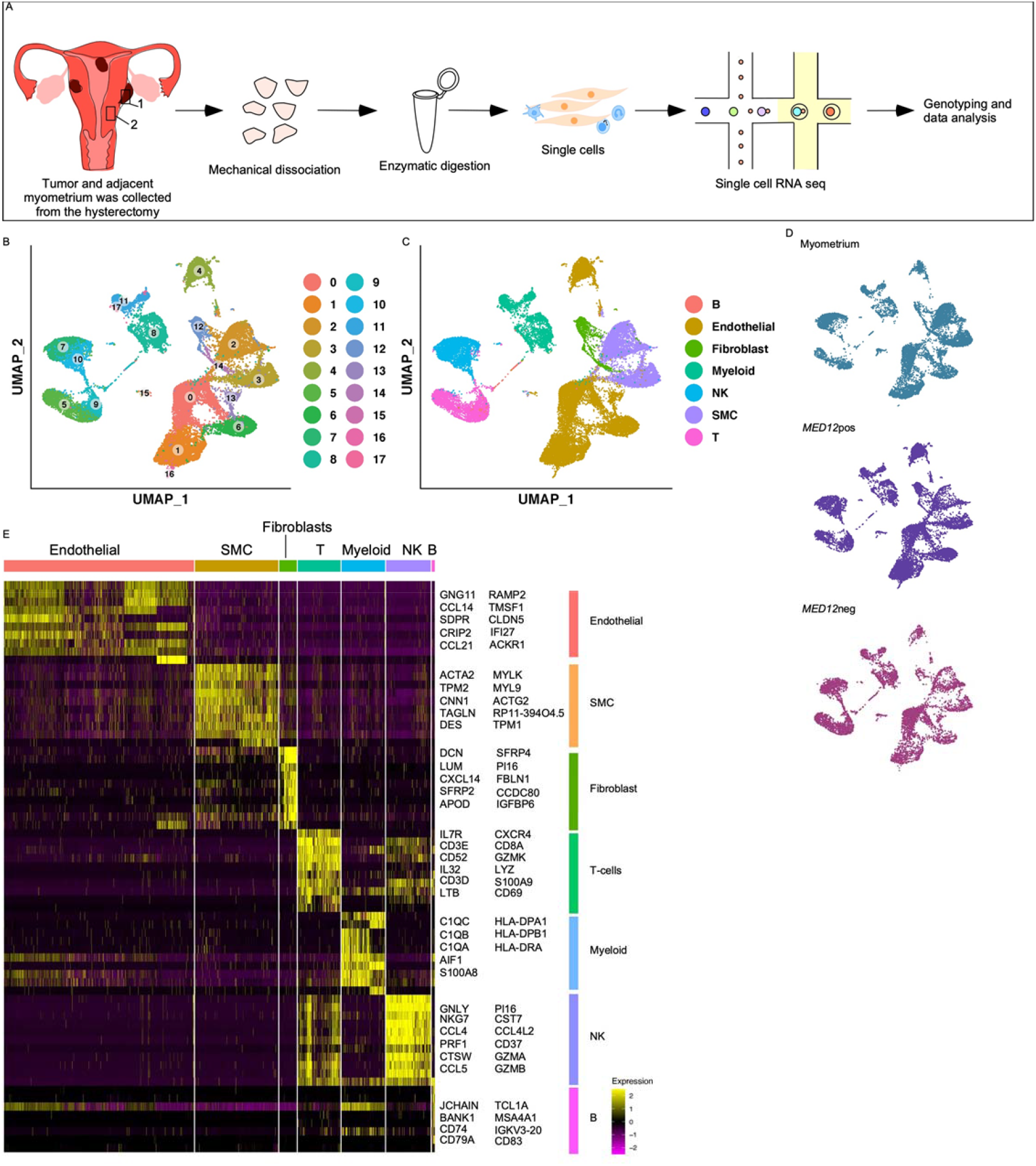
Single cell atlas of normal myometrium and leiomyomas. A) Overview of the dissociation protocol for both normal myometrium (n=5) and leiomyomas (n=8). B) Clustering of 34,435 high quality cells from *MED12* variant positive leiomyomas, *MED12* negative leiomyomas and normal myometrium. C) Cell lineages identified by the marker gene expression D) Annotation of the cell clusters per sample. E) Heatmap showing the top 10 genes used for cluster identification. Columns denote cells, rows denote genes.

### Characterization of the smooth muscle cells and fibroblast populations in normal myometrium

The cellular composition and molecular characterization of the normal myometrium are poorly understood. To examine the cellular heterogeneity in the myometrium, we first generated a UMAP from the myometrium dataset only. We identified the presence of 18 different cell clusters. These clusters included smooth muscle cells, fibroblast cells, endothelial cells, lymphatic endothelial cells, and immune cell populations (S Fig.2A).

To determine if there is an additional underlying heterogeneity in the SMC population of the normal myometrium, we extracted all the cells expressing *TAGLN*, *CNN1* and *SMA*, and clustered them at a higher resolution. These genes are known markers of SMC in uterus and other tissues^17,18^. Additional re-clustering of these cells revealed five different smooth muscle cell clusters in the normal myometrium (S Fig. 2B,F, Fig. 2E). All of the clusters were present in all patient samples, indicating reproducibility and absence of technical or batch effects (S Fig. 2C).

**Fig.2:**
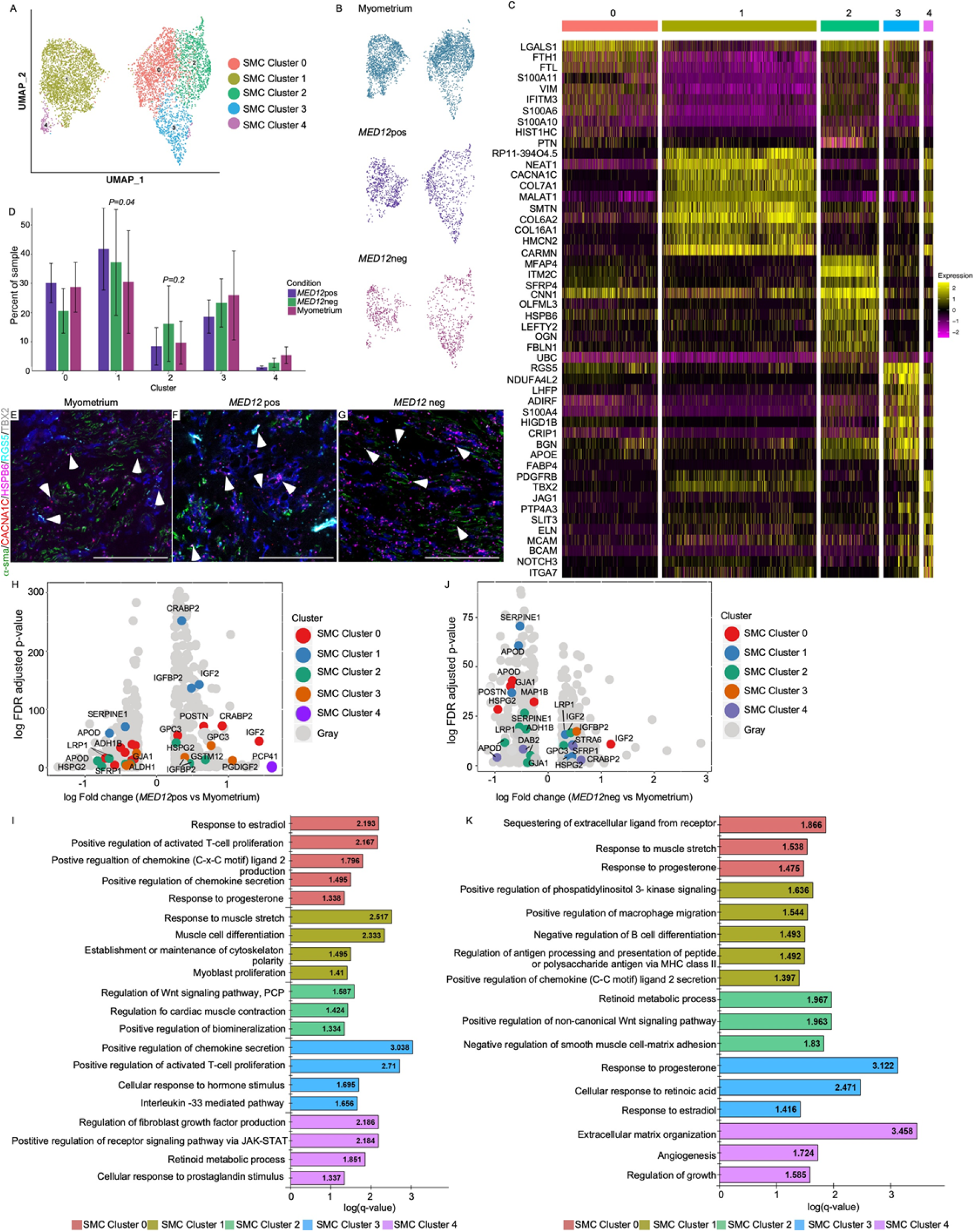
Intracellular heterogeneity and transcriptomic changes in the MED12 variant positive and MED12 variant negative smooth muscle cell clusters in comparison to the normal myometrium. A) UMAP showing the clusters of 6500 smooth muscle cells B) UMAP showing that all cell clusters are present in all three conditions-myometrium, *MED12* variant positive leiomyomas and *MED12* variant negative leiomyomas. C) Heatmap of the smooth muscle cell clusters. Colored bar on the top represents cluster number, Columns denote cells; rows denote genes. D) Cell proportion changes in the *MED12* variant positive and *MED12* variant negative leiomyomas in comparison to normal myometrium. Data represented here are mean ± s.e.m. *P* values were determined by Dirichlet-multinomial regression E-G) in situ images showing the validation of the smooth muscle cell clusters in *MED12* variant positive leiomyomas, *MED12* variant negative leiomyomas and normal myometrium. Scale bar is 100 μm. H) Volcano plots showing the transcriptomic changes in the smooth muscle cell clusters of *MED12* variant positive compared to the normal myometrium. I) GO analysis of DE genes showing at least 1.5 fold change in the *MED12* variant positive leiomyomas as compared to the normal myometrium J) Volcano plots showing the transcriptomic changes in the smooth muscle cell clusters of *MED12* variant negative compared to the normal myometrium K) GO analysis of DE genes showing at least 1.5 fold change in the *MED12* variant negative leiomyomas as compared to the normal myometrium.

To delineate the functional role of the SMC, we performed gene ontology (GO) enrichment analysis. Functional enrichment analysis with the biological function revealed that SMC Cluster 0 was specifically associated with intracellular sequestering of iron ion, cellular homeostasis, and response to stimulus. SMC Cluster 1 was defined by unique expression of the gene *CACNA1C* and was associated with muscle contraction, collagen fibril organization, extracellular matrix assembly and cellular response to TGF-β. SMC Cluster 2 was enriched in *HSPB6* and *MFAP4*. These cells were associated with glucocorticoid regulation, extracellular matrix (ECM) assembly and organization. SMC Cluster 3 was defined by the unique expression of genes *RGS5* and *HIGD1B*, which are specifically responsible for regulating the cellular response to TNF, regulation of endothelial cell proliferation. SMC Cluster 4 was enriched in the expression of *TBX2, PDGFRB, ELN, NOTCH3* and *MCAM* which are responsible for transcription by RNA polymerase II and showed genes upregulated for stem cell differentiation. A full list of the GO processes for all of the clusters is provided in the supplementary table (S Table 2). We performed *in situ* hybridization to validate the biological presence of these clusters, using the markers α.SMA^+^, *CACNA1C^-^*, *HSPB6^-^, RGS5^-^, TBX2^-^* (SMC Cluster 0), *CACNA1C* (SMC Cluster 1), *HSPB6* (SMC Cluster 2), *RGS5* (SMC Cluster 3), and *TBX2* (SMC Cluster 4) (Fig. 2E).

Our data analysis also identified two distinct cell subsets within fibroblast cell populations (S Fig. 2D,E). These fibroblasts differ from each other in the expression of *SFRP2, PLAC9, SERPINF1, S100A10* (S Fig. 2G). *In situ* hybridization confirmed the presence of two different fibroblast cell populations (Fig. 3E) We next investigated the biological function of these fibroblasts by performing gene ontology (GO) enrichment analysis. We found that the *WNT* regulated cluster (Fibro Cluster 0) was mainly involved in regulating fibroblast proliferation, ECM constituent secretion, and wound healing (S Table 3). Fibro Cluster 1 was linked to cellular component organization or biogenesis and regulation of mRNA splicing (S Table 3).

**Fig.3:**
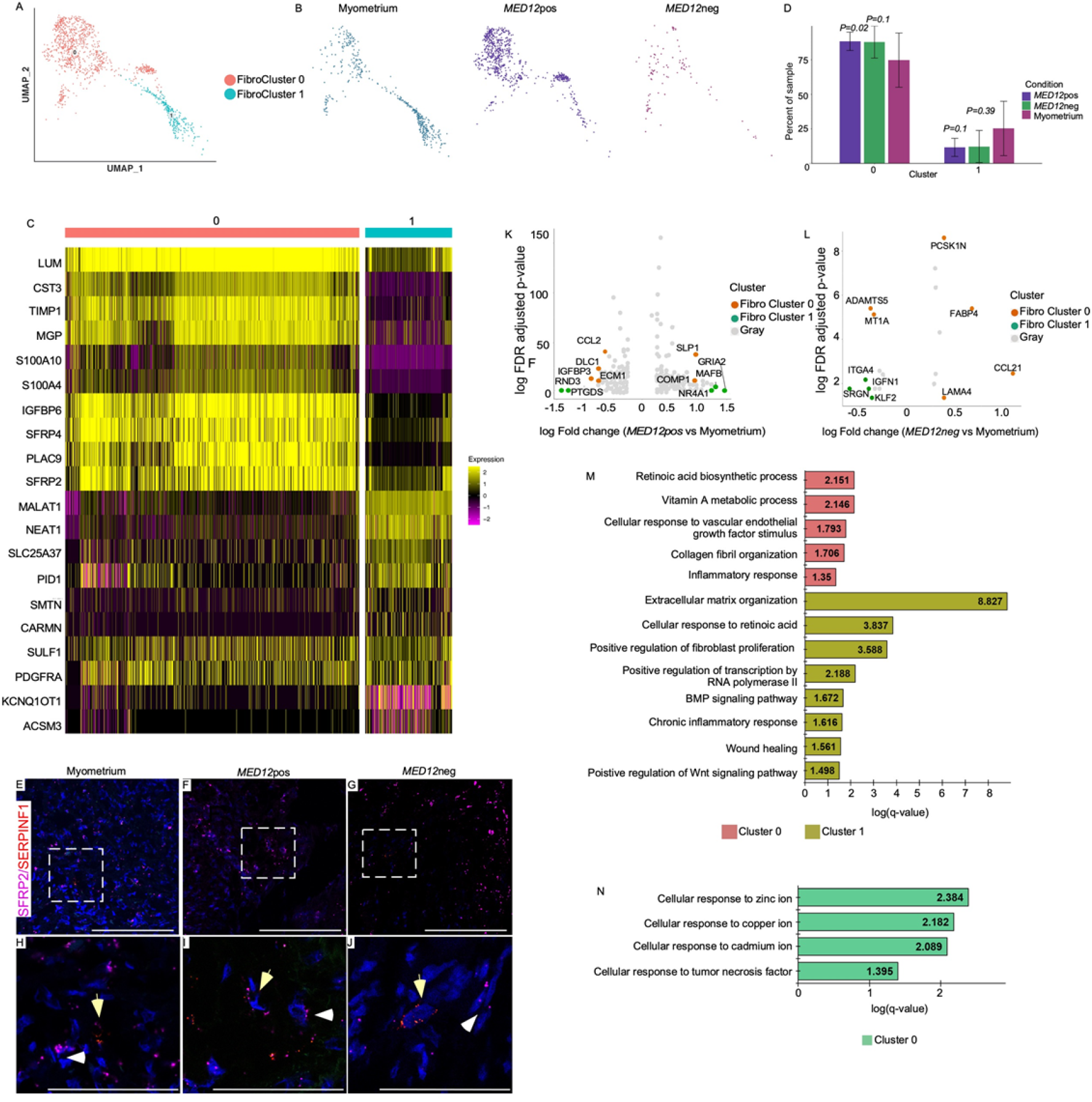
Intracellular heterogeneity and transcriptomic changes in the *MED12* variant positive and *MED12* variant negative fibroblast cell clusters in comparison to the normal myometrium. A) UMAP showing the clusters of 1291 fibroblast cells B) UMAP showing that all cell clusters are present in all three conditions-myometrium, *MED12* variant positive leiomyomas and *MED12* variant negative leiomyomas C) Heatmap of the fibroblast cell clusters. Colored bar on the top represents cluster number, Columns denote cells; rows denote genes. D) Cell proportion changes in the *MED12* variant positive and *MED12* variant negative leiomyomas in comparison to the normal myometrium. Data represented here are mean ± s.e.m. *P* values were determined by Dirichlet-multinomial regression E-J) *in-situ* images showing the validation of fibroblast populations *SFRP2* (Fibro Cluster 0; white arrowheads), and *SERPINF1* (Fibro Cluster 1; yellow arrowheads) in normal myometrium, *MED12* variant positive leiomyomas and *MED12* variant negative leiomyomas. Scale bar is 100 μm. K) Volcano plots showing the transcriptomic changes in the fibroblast cell clusters of *MED12* variant positive compared to the normal myometrium L) Volcano plots showing the transcriptomic changes in the fibroblast cell clusters of *MED12* variant negative compared to the normal myometrium M) GO analysis of DE genes showing at least 1.5 fold change in the *MED12* variant negative leiomyomas as compared to the normal myometrium. No DE genes were found in cluster 1 in comparison to the normal myometrium. N) GO analysis of DE genes showing at least 1.5-fold change in the *MED12* variant positive leiomyomas as compared to the normal myometrium

### Distinct smooth muscle cell populations and fibroblast populations expand in leiomyomas

To identify the differences in the cell composition between leiomyomas and normal myometrium, we generated a UMAP for the leiomyoma-only dataset. We identified the presence of same cell clusters as identified in the normal myometrium which were constituted of smooth muscle cells, fibroblast cells, endothelial cells, T-cells, B-cells, NK cells, and myeloid cell populations (S Fig. 3A,B).

Leiomyomas are tumors of smooth muscle cells that are thought to originate from a single mutant smooth muscle stem cell^11^. However, our data shows the presence of multiple cell types in leiomyomas which are similar to the normal myometrium (SFig. 2, 3). To understand if there is further intracellular heterogeneity in leiomyomas, and whether this heterogeneity is genotype dependent, we categorized leiomyoma samples as *MED12* variant positive and *MED12* variant negative based on the presence on *MED12* variant. Then, we isolated the smooth muscle, fibroblast, endothelial cells and immune cell populations from the integrated dataset and resolved them at higher resolution. Our data analysis revealed the presence of five different smooth muscle cell populations in both *MED12* variant positive and *MED12* variant negative leiomyomas (Fig. 2A-C). All of these smooth muscle cell clusters were also present in normal myometrium (SFig. 2A). However, we observed *CACNA1C* smooth muscle cell expansion (SMC Cluster 1) in *MED12* variant positive leiomyomas by 3-fold in comparison to the normal myometrium and *MED12* variant negative leiomyomas (*P value = 0.04*, three-fold change) (Fig. 2D-G). Whereas in *MED12* variant negative leiomyomas, SMC Cluster 2 (*HSPB6+*) expands compared to the normal myometrium (*P value* = 0.1, three-fold change Fig. 2D-G).

Next, we wanted to determine if similar heterogeneity exists in the fibroblast cell population in leiomyomas. We found presence of two distinct fibroblast subsets in all the three tissue samples, myometrium, *MED12* variant positive, and *MED12* variant negative leiomyomas (Fig. 3A-C). We observed a significant expansion of Fibro Cluster 0 in both *MED12* variant positive (*P value* = 0.02, 1.14-fold change) and *MED12* variant negative (*P value* = 0.1, 1.14-fold change) leiomyomas compared to normal myometrium (Fig. 3D). This cluster seems to be regulated by the *WNT* signaling pathway (Fig. 3C). *In situ* hybridization for *SFRP2* confirmed the expansion of the *WNT* regulated fibroblasts in the *MED12* variant positive and *MED12* variant negative fibroblasts (Fig. 3E-J). Together, these datasets show the presence of a heterogenous smooth muscle cell and fibroblast cell populations in both *MED12* variant positive and *MED12* variant negative leiomyomas.

### Remodeling of the smooth muscle and fibroblast cell populations in leiomyomas compared to the myometrium

Our previous study has shown that mutation in *MED12* is sufficient to cause leiomyoma formation^19^. In this study we found that normal myometrium, and leiomyomas have similar cellular composition (Fig. 1B-D, S Fig. 2A, S Fig. 3A). We hypothesized that transcriptomic changes in normal myometrium due to the acquired genetic mutations are responsible for the leiomyoma formation.

To determine that, we compared the transcriptomic changes in smooth muscle cell subsets of *MED12* variant positive leiomyomas, and *MED12* variant negative leiomyomas to the normal myometrium. Gene ontology analysis of the *MED12* variant positive leiomyomas revealed enrichment for genes involved in the regulation of fibroblast proliferation, smooth muscle cell proliferation, inflammatory response, extracellular matrix organization, cellular response to hormones and proliferation of activated T-cells. These genes were found across the smooth muscle cell cluster in *MED12* variant positive leiomyomas compared to myometrium (Fig, 2H-K, S Table 4). We found significant expansion of the SMC Cluster 1 (*CACNA1C*) in *MED12* variant positive leiomyomas. These cells are enriched across genes regulating myoblast proliferation, response to muscle stretch, muscle cell differentiation, regulation of fibroblast growth factor production, and establishment and maintenance of cytoskeleton polarity (Fig. 2H,I, S table 4).

We found that SMC Cluster 2 was most enriched in the *MED12* variant negative leiomyomas in comparison to the normal myometrium (Fig 2D). Further analysis of the transcriptomic changes in the smooth muscle cells of the *MED12* variant negative leiomyomas compared to the normal myometrium revealed that in SMC Cluster 2, genes involved in the biological processes such as retinoid metabolic process (*GPC3*, *STRA6*, *ADH1B, CRABP2, HSPG2, LRP1*), positive regulation of non-canonical *WNT* signaling pathway (*GPC3, DAB2, SFRP1*) and negative regulation of smooth muscle cell-matrix adhesion (*SERPINE1, APOD*) were enriched (Fig 2J,K, S Table 4). The full list of other transcriptomic changes in the smooth muscle clusters are provided in S table 4.

Previous studies suggest that one of the reasons for the size difference between *MED12* variant positive and *MED12* variant negative leiomyomas is due to differences in their cellular composition. They have suggested that in *MED12* variant positive leiomyomas both smooth muscle cells and fibroblast cell populations secrete collagen^**10**^. So, we decided to compare the transcriptional changes in *MED12* variant positive and *MED12* variant negative leiomyomas’ fibroblast subset to that of the myometrium. We found upregulation of the Vitamin A metabolic process, retinoic acid biosynthesis process, cellular response to the vascular endothelial growth response, collagen fibril formation and inflammatory response in *MED12* variant positive leiomyoma’s Fibro Cluster 0 (Fig. 3K,M). This fibroblast population is regulated by *WNT* signaling in normal myometrium and expands in the *MED12* variant positive leiomyomas (Fig. 3D). While *MED12* variant positive Fibro Cluster 1 had genes induced for chronic inflammatory response, wound healing, BMP signaling, ECM organization and positive regulation of the transcription by RNA polymerase II (Fig. 3L,N). Interestingly, similar to *MED12* variant positive leiomyomas Fibro Cluster 0 also expands in *MED12* variant negative leiomyomas. While we observed significant transcriptional changes in both fibroblast cell clusters 0 and 1 in *MED12* variant positive leiomyomas as compared to normal myometrium, only *WNT* regulated fibroblasts (Fibro Cluster 0) showed transcriptional differences in *MED12* variant negative leiomyomas compared to the normal myometrium (Fig. 3L,N, Supplementary Table 5). We did not observe any significant transcriptional differences in Fibro cluster 1 between normal myometrium and *MED12* variant negative leiomyomas (Fig. 3L,N). Collectively, these data suggest that smooth muscle cells and fibroblast cells expand and remodel differently according to the leiomyoma’s genotype.

### Novel lymphatic endothelial cells inhabit leiomyomas

Unsupervised clustering of endothelial cells revealed the presence of nine clusters of endothelial cells (Fig. 4A-C). We observed that Endo cluster 8 was exclusively expressed in both *MED12* positive and *MED12* negative leiomyomas but not in normal myometrium (Fig. 4B,D). The Endo cluster 8 cells were present in all patient leiomyoma samples (S Fig. 4A). Based on differential gene expression, we identified these cells as lymphatic endothelial cells (Fig. 4C). Violin plots and gene ontology analysis of Endo Cluster 8 further confirmed the lymphatic endothelial cell fate of this cell cluster (S Fig. 4B,C). In addition to the lymphatic endothelial cell fate commitment genes (*PDPN*, *RELN*), Endo Cluster 8 also showed gene expression connected to DNA replication, mitosis and regulation of cytoskeleton organization (*CCL21, STMN1, FSCN1, PROX1, TPX2, MAP1B, PLK1, S100A10, NEK2*) (S Fig. 4B). These cells showed upregulation of genes associated with inflammation such as *PPFIBP1, SEPP1, PTX3, SPHK1* (S Table 6).

**Fig.4:**
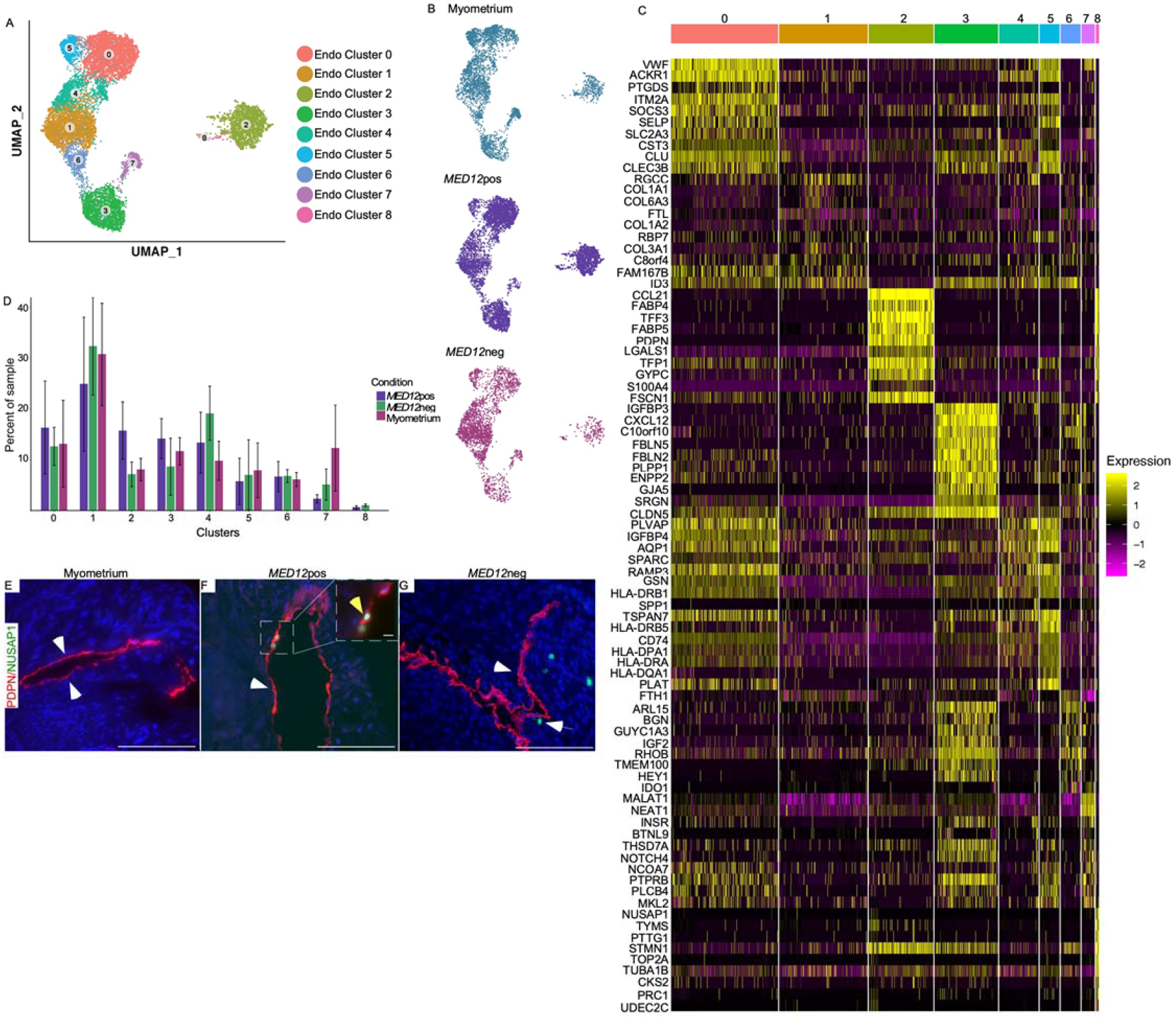
Novel lymphatic endothelial cells are present in leiomyomas. A) UMAP showing the clusters of 14742 endothelial cells B) UMAP showing the cluster annotation per condition-myometrium, *MED12* variant positive leiomyomas and *MED12* variant negative leiomyomas C) Heatmap of endothelial cell clusters. Colored bar on the top represents cluster number, Columns denote cells; rows denote genes. D) Cell proportion changes in the *MED12* variant positive and *MED12* variant negative leiomyomas in comparison to the normal myometrium. Data represented here are mean ± s.e.m. *P* values were determined by Dirichlet-multinomial regression E) Immunostaining for PDPN (white arrowheads) and *NUSAP1* (yellow arrowheads) showing presence of lymphatic endothelial cell clusters in *MED12* variant positive leiomyomas, *MED12* variant negative leiomyomas and normal myometrium. Scale bar is 100 um.

Interestingly, Endo cluster 2, also identified as a lymphatic endothelial cell cluster, was present in all three sample types, normal myometrium, *MED12* variant positive and *MED12* variant negative leiomyomas. Endo Cluster 8 was differentiated from Endo Cluster 2 by the expression of Nucleolar-spindle-associated protein (*NUSAP1*), which is an important regulator of mitosis^20^. We confirmed the leiomyoma specific expression of Endo Cluster 8 by immunostaining (Fig. 4E-G). Studies have shown that inflammation is responsible for lymphangiogenesis in different conditions and is responsible for the immune response in these respective conditions^21^. To confirm if the cells in Endo Cluster 8 originated from Endo Cluster 2 in response to the inflammation, we performed pseudotime analysis. Our analysis revealed the expression of Ki67, a proliferation marker in Endo Cluster 8 (S.Fig. 5A,B). Therefore, these data indicate that these cells may have originated from Endo Cluster 2 in leiomyomas in response to the inflammation.

### Immune cell infiltration differs by the genotype of leiomyomas

Next, we wanted to analyze the immune profile of the normal myometrium and compare it to the leiomyomas. Our data showed the presence of two subsets of natural killer (NK) cells, B-cells, dendritic cells, T-cells, CD8+ve T-cells, CD14 monocytes, FCER3A+ monocytes, and two subsets of macrophages in the normal myometrium (Fig. 5A-C). These results are in agreement with the previously reported immune cell populations in myometrium^22^. Missing cell types in our data that were previously reported to be present in the myometrium are mast cells and neutrophils ^22–24^.

**Fig.5:**
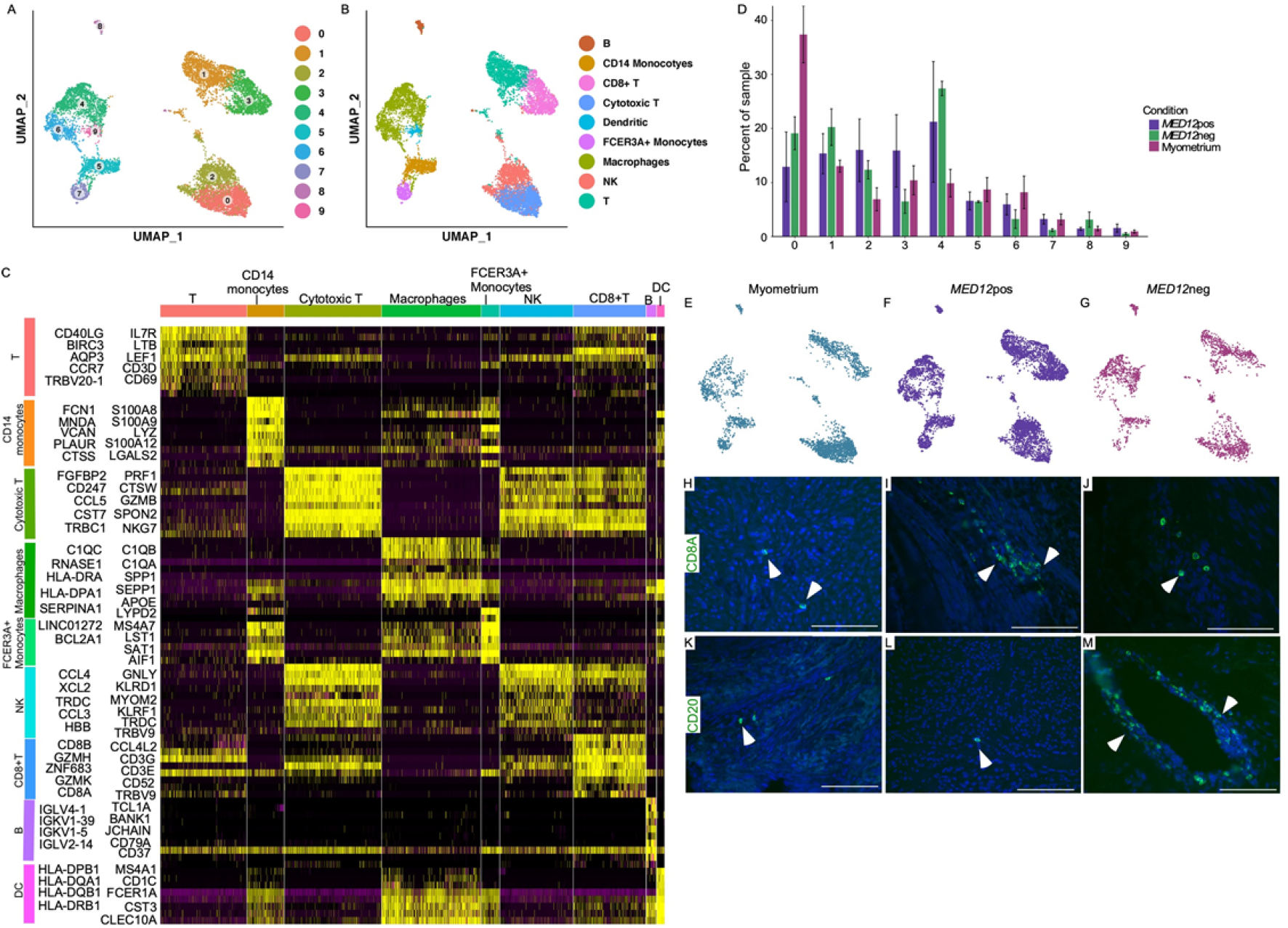
Immune cell infiltration is dependent on genotype of the uterine leiomyoma. A) UMAP showing the immune cell clusters. B) Annotation of the UMAP clusters. C) Heatmap of endothelial cell clusters. Colored bar on the top represents cluster number. Columns denote cells; rows denote genes. D) Cell proportion changes in the *MED12* variant positive and *MED12* variant negative leiomyomas in comparison to the normal myometrium. Data represented here are mean ± s.e.m. *P* values were determined by Dirichlet-multinomial regression E-G) UMAP showing the cluster annotation per condition-myometrium, *MED12* variant positive leiomyomas and *MED12* variant negative leiomyomas respectively. H-J) CD8a (white arrowheads) showing increase infiltration of T-cells–in *MED12* variant positive leiomyomas, as compared to normal myometrium and *MED12* variant negative leiomyomas. K-M) CD-20 staining (white arrowheads) showing increased infiltration of B-cells in *MED12* variant negative leiomyomas compared to normal myometrium and *MED12* variant positive leiomyomas. Scale bar is 100 um.

We found an increase in infiltration of T-cells in the *MED12* variant positive leiomyomas (Fig. 5D, H-J). These cells express cytokines and chemoattractant for the recruitment of CD8+ T-cells which have increased cytotoxic activity, indicated by upregulation of *GNLY, RHOB*, and genes associated with the antigen recognition (*TRBV7-9, TRBV9, TRAV38-2DV, TYROBP*) (S Fig. 6C; S Table 7). GO enrichment analysis for the biological processes showed upregulation of genes associated with the adaptive immune response and regulation of the immune response (S Fig. 6A). We also found expansion of NK cells expressing *CCL21* and *CD96* in the *MED12* variant positive leiomyomas. These cells showed upregulation of IL-6 secretion, a proinflammatory cytokine and genes responsible for the regulation of the inflammatory response to the antigen stimulation (S Table. 7). Expansion of T-cells, NK cells, and dendritic cells which participate in the adaptive immune response and lymphatic endothelial cells in the *MED12* variant positive leiomyomas further suggest that activation of the adaptive immune response plays a role in the management of the *MED12* variant positive leiomyomas.

In contrast, we observed expansion of macrophages and B-cells in the *MED12* variant negative leiomyomas (Fig. 5K-M). Macrophages were enriched for ontology terms relevant to regulation of B-cell differentiation and regulation of angiogenesis in addition to known macrophage-functions such as antigen processing (S Fig. 6B). B-cells showed upregulation of the genes *IGLV3-11, IGLV3-21*, and *FCER1G1* which are responsible for antigen recognition and affinity maturation (S Fig. 6D, S Table 7). Tissue immunostaining confirmed the expansion of macrophages and B-cells in the *MED12* variant negative leiomyomas compared to *MED12* variant positive leiomyomas and normal myometrium (Fig. 5D, K-M, S Fig. 6E-G). These data highlight that immune cell infiltration and the immune response to leiomyomas is dependent on the genotype.

### *MED12* leiomyoma cell types are not monoclonal

In normal female cells, monoallelic expression of X-chromosome genes is observed due to random inactivation of one of the two X-chromosomes for dosage compensation^25^. This principal is widely used to study the clonality in tumors ^26^. Based on this phenomenon, studies utilizing random inactivation of X-linked gene human androgen receptor (*HUMARA*) and inactive glycose-6-phosphate dehydrogenase (*G6PD*) isoform expression have suggested that uterine leiomyomas are monoclonal tumors of smooth muscle cells of the uterus ^27,28^. However, our single cell sequencing data indicates the presence of cellular heterogeneity in the smooth muscle cell populations, fibroblast populations, and endothelial cell populations in the leiomyomas (Fig. 1B, C). Moreover, *MED12* is located on the X-chromosome and is subject to X-chromosome inactivation. We hypothesized that if leiomyomas are truly monoclonal and derive from a so called “stem cell”, then different cell types identified in leiomyomas should only express the *MED12* mutant variant. Single cell RNA sequencing has an advantage over the previously used techniques to determine the clonality of the tumor because of the ability to capture variants in individual cells and interrogate at higher resolution.

We therefore performed variant analysis of the single-cell sequencing datasets of *MED12* variant positive leiomyomas using Integrative Genomics Viewer (IGV). IGV analysis revealed expression of both the mutant variants (G44T, G44C) and the wild type *MED12* allele in the *MED12* variant positive leiomyomas (Fig. 6A). To determine which cells express *MED12* variant, we generated a UMAP showing smooth muscle cells, fibroblasts and endothelial cells from three *MED12* positive fibroid samples (Fig. 6B). We generated the UMAP from only these cells to determine if expression of wild type *MED12* is a result of immune cell infiltration in the tumor. We then identified the *MED12* variants in the single-cell RNA sequencing data by utilizing the dsc-pileup software developed from previously published tool, demuxlet^15,16^. We plotted the *MED12* variants in the UMAP generated using the smooth muscle cell, fibroblast, and endothelial cell populations of the *MED12* leiomyomas. We found expression of *MED12* variants (*MED12* G>N - 40%, *MED12* wild type allele-60%) in all three cell populations (Fig. 6C). The expression of the *MED12* variant allele and wild type allele varies from patient to patient with some patients expressing roughly 80% of the *MED12* variant allele while others expressed only 20% of the *MED12* variant allele (Fig. 6A). These results indicate that *MED12* variants present in a limited number of myometrium cells might be sufficient to cause leiomyomas. Upon plotting the wild type *MED12* allele of these cells, we found the presence of the wild type *MED12* in all clusters of smooth muscle cells, fibroblasts and endothelial cell (Fig. 6C). For the first time, our data indicates that *MED12* variant positive leiomyomas might not be monoclonal in nature.

**Fig.6:**
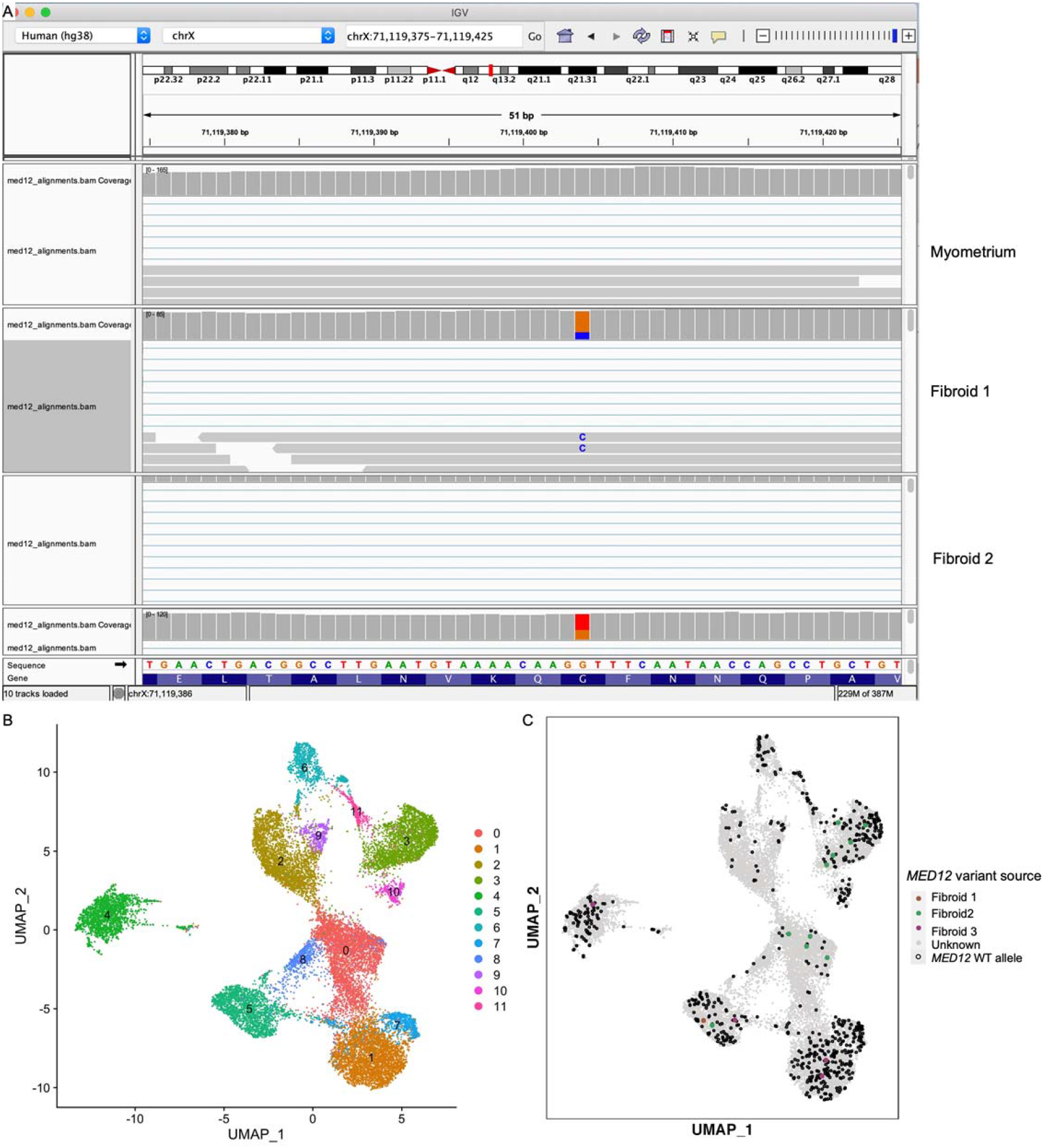
*MED12* variant positive uterine leiomyomas are not monoclonal. A) IGV analysis showing presence of both the variant and wild type codon at c131 in the *MED12* variant positive leiomyomas. B) UMAP of the mesenchymal cell populations in the leiomyomas. Clusters 0, 1, 5, 7, 8 are endothelial cells, Cluster 2, 3 and 9 are smooth muscle cells, clusters 11 and 6 are fibroblasts, clusters 4 and 10 are lymphatic endothelial cells. C) UMAP showing the presence of the mutant *MED12* variant (colored dots) in all smooth muscle cells, fibroblasts and endothelial cells and the wild type *MED12* (black dots).

## Discussion

Uterine leiomyomas are thought to originate from SMC in myometrium. However, recent studies have reported presence of fibroblasts and endothelial cells in uterine leiomyomas in addition to SMC populations^10,12^, thus suggesting presence of cellular heterogeneity in leiomyomas. We applied single cell RNA sequencing technique to create a cell atlas for the normal myometrium and the leiomyomas. Our data shows that at least 18 different cell types compose myometrium, including different subtypes of smooth muscle cells, fibroblast, vascular endothelial cells, lymphatic endothelial cells and immune cells. Other than novel lymphatic endothelial cell population exclusive to leiomyomas, myometrium and leiomyomas share similar cell composition. It is therefore likely that transcriptomic differences account for leiomyomagenesis.

Previous studies have established the differences in the transcriptomic profile between the normal myometrium and leiomyomas^29,30^. These studies have consistently identified differential expression of genes regulating ECM metabolism (*COL12A1, COL6A3, FN1, ADAM19*), hormone responsiveness (*PRLHR, EGFR, CYP26A1, EGR1*), and muscle physiology (*RGS2*, *CACNA1C, MYH2*)^29,30^. We analyzed the transcriptional changes in leiomyomas and myometrium at the single cell resolution. Similar to the previous reports we found upregulation of genes involved in ECM metabolism (*COL7A1, COL6A2, COL16A1*), hormone responsive genes (*PTN*, *FOSB*, *IGFBP2*, *FOS*), and muscle organization and differentiation (*TTN*, *LMNA*, *RGS2*) in leiomyomas compared to the normal myometrium. We also examined the effect of underlying leiomyoma genotype on transcriptomic changes. Previous studies have shown that missense variants in *MED12* exon 2 associate with 70% of leiomyomas^4–6^. We compared the transcriptomic changes at single-cell level among *MED12* variant positive and *MED12* variant negative leiomyomas against the myometrium. We found that both SMC and fibroblast cells show upregulation of genes associated with ECM metabolism in *MED12* variant positive leiomyomas, while in *MED12* variant negative leiomyomas, only SMC showed such upregulation. Our results are consistent with previous report that both SMC and fibroblasts are responsible for ECM production in *MED12* variant positive leiomyomas^10^. Although ECM accumulation is a characteristic feature of leiomyomas there is strikingly increased ECM accumulation in *MED12* variant positive leiomyomas compared to *MED12* variant negative leiomyomas^10^. It is possible that these transcriptional differences explain increased accumulation of ECM in *MED12* variant positive leiomyomas compared to *MED12* variant negative leiomyomas.

Previous studies using human tissue and transgenic mouse models have implicated the role of Wnt/β-catenin, REST-GPR-10, and mTOR signaling pathways in leiomyoma pathology^29,31–34^. In agreement to these studies, our human scRNA seq data here shows dysregulation of WNT signaling, retinoic acid, PI3K, and JAK-STAT signaling pathways in different cell clusters of SMC and fibroblast populations in both *MED12* variant positive and *MED12* variant negative leiomyomas. However, we found an individual SMC or fibroblast cell cluster can have dysregulation of multiple signaling pathways in both *MED12* variant positive and *MED12* variant negative leiomyomas. These results indicate that a specific signaling pathway might not be responsible for leiomyoma formation. We have previously shown that common nonsynonymous *MED12* exon 2 variant (c.131G>A) causes leiomyomagenesis in a mouse model ^19^. Using our *MED12* mouse model, we found dysregulation of *Wnt* signaling pathway, *Ras* signaling, and mTOR signaling pathways in leiomyomas^19^The involvement of multiple pathways is likely due to *MED12*, which plays a known role as part of the mediator complex that regulates global RNA polymerase II dependent transcription and *MED12* variant is likely to have global effect^5^. *MED12* variant negative leiomyomas have a highly heterogenous genetic landscape and many of these leiomyomas carry *HMGA2* rearrangements^5,7^. *HMGA2* belongs to a non-histone chromosomal high-mobility group (HMG) protein family and acts as a transcriptional regulating factor associated with multiple pathologies^35,36^.

Uterine leiomyomas are hormone responsive in nature^37^. Previous studies have shown that leiomyomas increase in size with the presence of estrogen and progesterone^11,19^. Leiomyomas with *HMGA2* rearrangements and *MED12* mutations show increased proliferation of SMC in the presence of estrogen and progesterone, whereas fibroblast population proliferates solely in response to estrogen^10^. Our study shows that all SMC clusters in *MED12* variant positive and *MED12* variant negative leiomyomas have upregulation of both estrogen and progesterone responsive genes However, we did not observe an enrichment of estrogen or progesterone responsive genes in leiomyoma fibroblast cell clusters (Fig 2I,J, 3M,N, S Table 4,5). Our results indicate that SMCs may be more responsive to ovarian hormones than fibroblasts. It remains to be studied if the fibroblast cells carrying *MED12* variant behave like SMC and show increased responsiveness to ovarian hormones or if they interact with SMC in paracrine fashion.

Lymphatic endothelial cells were previously reported in myometrium but not in leiomyomas^38,39^. In this study, we found that while normal myometrium has only one lymphatic endothelial cell population, there are two distinct populations of lymphatic endothelial cells in both *MED12* variant positive and *MED12* variant negative leiomyomas. The lymphatic endothelial cells are known to facilitate the recruitment of immune cells to tissues^40^. Previous studies have reported variable predominance of T-cells and B-cells in leiomyomas^41^. Unexpectedly, we found that the differences in immune cell infiltration in leiomyomas are genotype dependent. While the *MED12* variant positive leiomyomas showed increased infiltration of T-cells and NK cells. *MED12* variant negative leiomyoma show predominance of B-cells and macrophages suggesting activation of humoral immune response. The increased infiltration of the immune cells in leiomyomas compared to the normal myometrium can be explained by the presence of additional lymphatic endothelial cells. Why the immune cell infiltration in the leiomyomas differs with respect to their genotype and functional significance needs to be explored in future studies. Collectively, these findings demonstrate underlying cellular heterogeneity in both leiomyomas and myometrium. Our data also explains that previously reported immune cell population infiltration variability in leiomyomas is genotype dependent.

Based on studies utilizing HUMARA assay and inactivation of glucose-6-phosphate dehydrogenase isoform expression, uterine leiomyomas are widely accepted to be monoclonal in nature^27,28^. These assays are widely used to study the clonality in different tumors and tissues^42^. However, recent studies have questioned the accuracy of these assays partly affected by inconsistent methylation of X-chromosome genes^43,44^. *MED12* mutation is responsible for leiomyoma formation in 70% of women^4^. *MED12* is located on the X-chromosome and because of X chromosome inactivation either the variant or the wild type *MED12* allele will be expressedl^45^. It has previously been shown that *MED12* variant positive leiomyomas express the variant allele^4,5^. If leiomyomas were monoclonal and originated from a single *MED12* variant carrying stem cell, then all of the cells present in a leiomyoma would be expected to carry the same *MED12* variant. Our analysis of the data, exploiting the unique genotype of *MED12* variant positive leiomyomas, shows that uterine leiomyomas cellular moiety is not monoclonal in nature. We found that *MED12* variant positive leiomyomas were composed of a mixture of cells expressing both *MED12* variant allele as well as *MED12* wild type allele. Other studies using alternative techniques support our findings. FACS sorting of leiomyomas into SMC and fibroblast populations revealed that *MED12* variant positive leiomyomas are composed of SMC carrying *MED12* variant allele while the fibroblasts cells in leiomyomas do not carry the *MED12* variant^10^. The study concluded that the leiomyoma causative mutations are present in SMC only. We, however, found that leiomyomas are a mixture of cells expressing both *MED12* variant and wild type allele in multiple cell types examined including SMC, and endothelial cells. Although our results show that leiomyoma cellular moiety is not monoclonal, the origin of leiomyomas remains uncertain.

In conclusion, the single cell atlas of human myometrium and leiomyomas show cellular heterogeneity and complexity that is genotype dependent. The main differences include novel, leiomyoma specific lymphatic endothelial cell type, genotype dependent immune infiltration and transcriptomic changes that may account for hormone responsiveness and ECM accumulation. Moreover, our studies show that leiomyoma cell moiety is not monoclonal in nature, which should be a major consideration when designing future therapeutics against leiomyomas.

## Code availability

The datasets generated and analyzed in the study are available in the NCBI Gene Expression Omnibus (GEO) and Sequence Read Archive (SRA) and can be accessed upon request. All custom scripts can be accessed upon request to the Lead Contacts.

## Contributions

JG and AR conceived the study, JG designed and performed the experiments, data analysis and interpretation. AR supervised the study, designed experiments, and performed data analysis. JR performed computational analysis. JJW helped with the pathological examination of the tissues used in the study. S.B and D.C helped with data analysis. JG and AR wrote the manuscript with input from all the authors.

## Acknowledgments

We would like to thank Dr. Rohit K. Gupta for helping us with the sample preparation protocol. Meghana Sukhthankar for helping with the sample preparation. Dr. Katja Rust and Tania Moody for discussions about the data analysis. Matthew Tae for help with GO analysis. We would like to thank all the Rajkovic lab members for the critical reading of the document. This work was supported by funding from National Institute of Child Health and Human Development (5P50HD098580).

