## Supplementary files for "Single Cell atlas of uterine myometrium and leiomyomas reveals diverse and novel cell types of non-monoclonal origin"

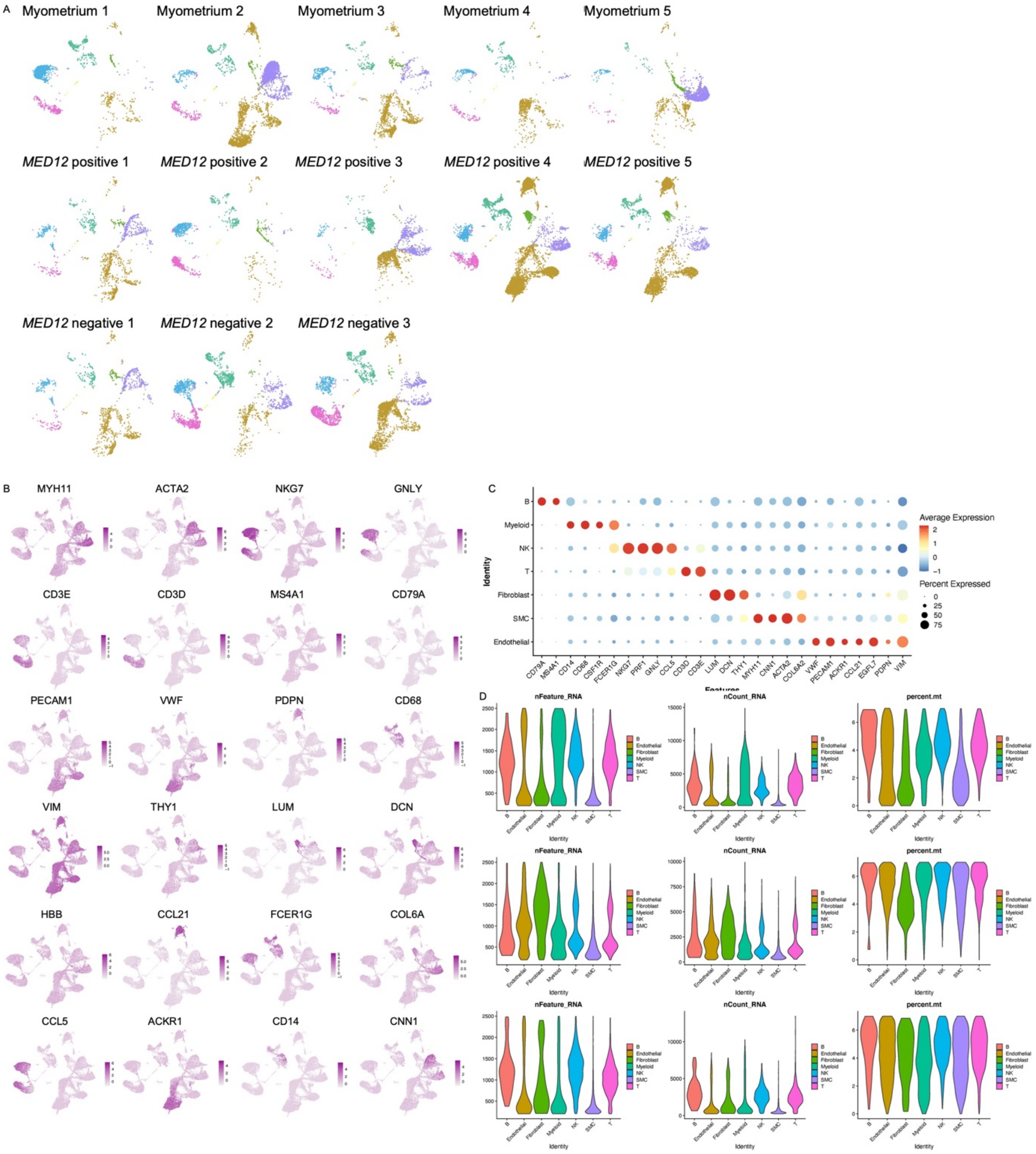
A
Myometrium 1
Myometrium 2
Myometrium 3
Myometrium 4
Myometrium 5
MED12 positive 1
MED12 positive 2
MED12 positive 3
MED12 positive 4
MED12 positive 5
MED12 negative 1
MED12 negative 2
MED12 negative 3
B
MYH11
ACTA2
NKG7
GNLY
CD3E
CD3D
MS4A1
CD79A
PECAM1
VWF
PDPN
CD68
VIM
THY1
LUM
DCN
HBB
CCL21
FCER1G
COL6A
CCL5
ACKR1
CD14
CNN1
C
Identity
B
Myeloid
NK
T
Fibroblast
SMC
Endothelial
CD79A
MS4A1
CD14
CD68
CSF1R
FCER1G
NKG7
PRF1
GNLY
CCL5
CD3D
CD3E
LUM
DCN
THY1
MYH11
CNN1
ACTA2
COL6A2
VWF
PECAM1
ACKR1
CCL21
EGFL7
PDPN
VIM
Features
Average Expression
Percent Expressed
D
nFeature_RNA
nCount_RNA
percent.mt
Identity
B
Endothelial
Fibroblast
Myeloid
NK
SMC
T

**S Fig. 1: Quality control and annotations of the myometrium cells** A) UMAP showing the cell types were present in all individual patient samples including myometrium, *MED12* variant positive leiomyoma and *MED12* variant negative leiomyomas B) Feature plots showing the expression of the lineage markers of the cell types (deep purple: high; light purple: low) C) Dot plots showing the lineage signatures per cell type. The dot size indicates the percentage of expression and the color of the dot is indicative of average expression of the gene (red: high, blue: low) D) Violin plots of the number of genes, number of total counts and percentage mitochondrial genes expressed in the across 34,435 cells in myometrium, *MED12* variant positive and *MED12* variant negative leiomyoma samples.

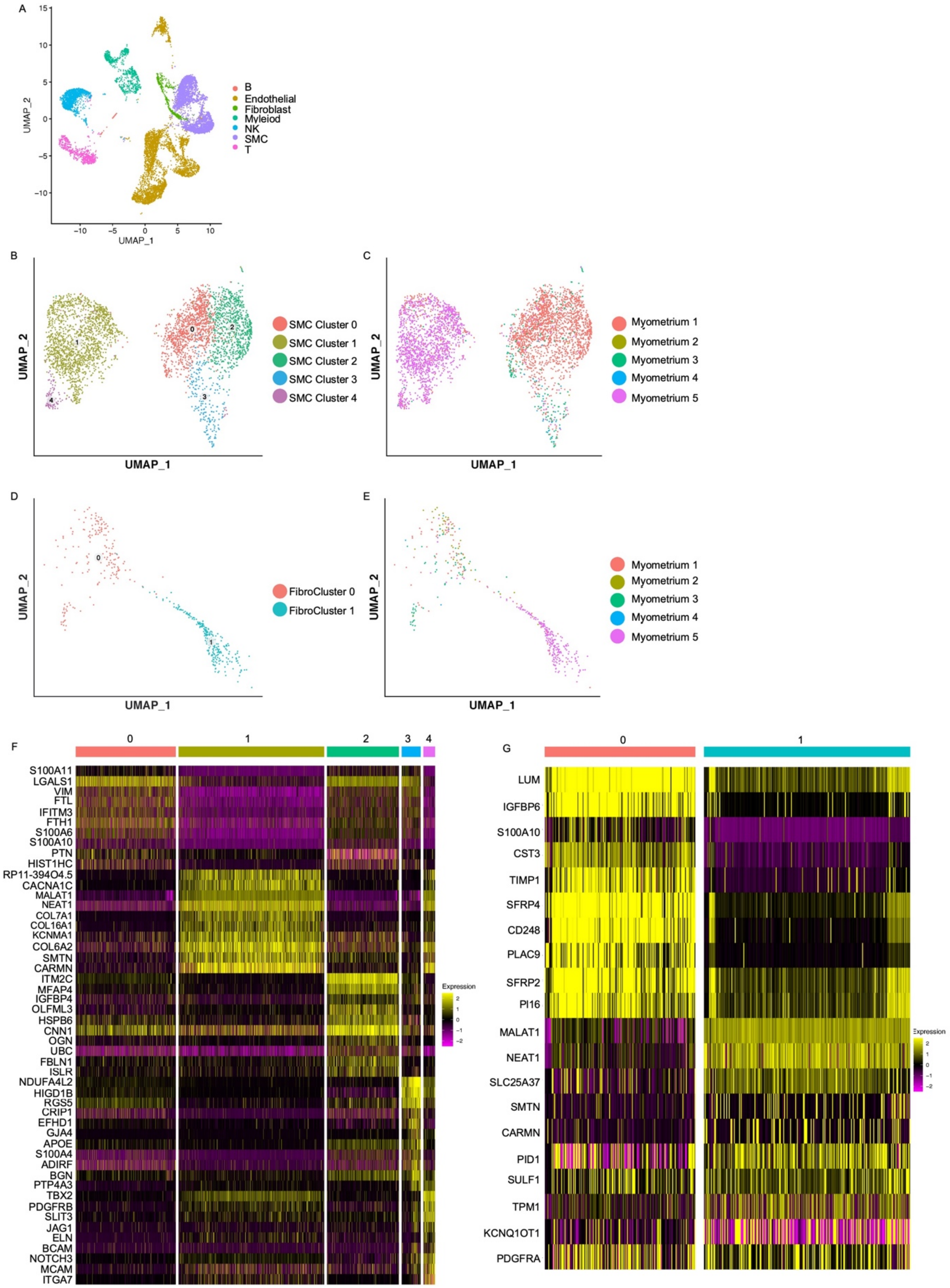
A
UMAP_1
UMAP_2
B
Endothelial
Fibroblast
Myleiod
NK
SMC
T
B
SMC Cluster 0
SMC Cluster 1
SMC Cluster 2
SMC Cluster 3
SMC Cluster 4
C
Myometrium 1
Myometrium 2
Myometrium 3
Myometrium 4
Myometrium 5
D
FibroCluster 0
FibroCluster 1
E
F
S100A11
LGALS1
VIM
FTL
IFITM3
FTH1
S100A6
S100A10
PTN
HIST1HC
RP11-394O4.5
CACNA1C
MALAT1
NEAT1
COL7A1
COL16A1
KCNMA1
COL6A2
SMTN
CARMN
ITM2C
MFAP4
IGFBP4
OLFML3
HSPB6
CNN1
OGN
UBC
FBLN1
ISLR
NDUFA4L2
HIGD1B
RGS5
CRIP1
EFHD1
GJA4
APOE
S100A4
ADIRF
BGN
PTP4A3
TBX2
PDGFRB
SLIT3
JAG1
ELN
BCAM
NOTCH3
MCAM
ITGA7
Expression
G
LUM
IGFBP6
S100A10
CST3
TIMP1
SFRP4
CD248
PLAC9
SFRP2
PI16
MALAT1
NEAT1
SLC25A37
SMTN
CARMN
PID1
SULF1
TPM1
KCNQ1OT1
PDGFRA
Expression

**S Fig. 2: Intracellular heterogeneity in the normal myometrium.** A) UMAP showing all cell clusters present in the myometrium dataset B) UMAP showing the clusters of smooth muscle cells C) UMAP showing the smooth muscle cell clusters per individual patient sample D) UMAP showing the clusters of fibroblasts E) UMAP showing the fibroblast cell clusters per individual patient sample F) Heatmap of the smooth muscle cell clusters. Colored bar on the top represents cluster number. Columns denote cells; rows denote genes. G) Heatmap of the fibroblast cell clusters. Colored bar on the top represents cluster number. Columns denote cells; rows denote genes.

A

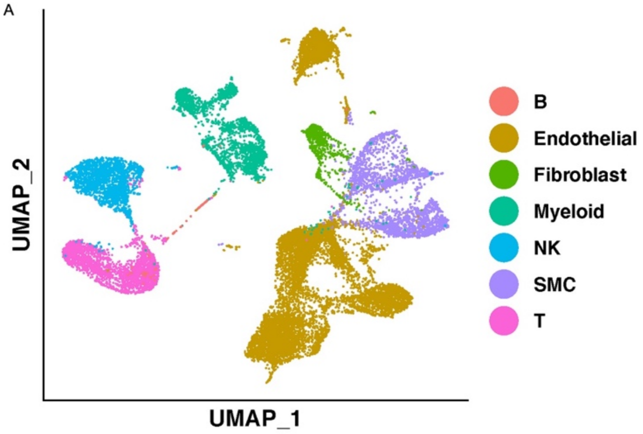
UMAP_2
UMAP_1
B
Endothelial
Fibroblast
Myeloid
NK
SMC
T

B

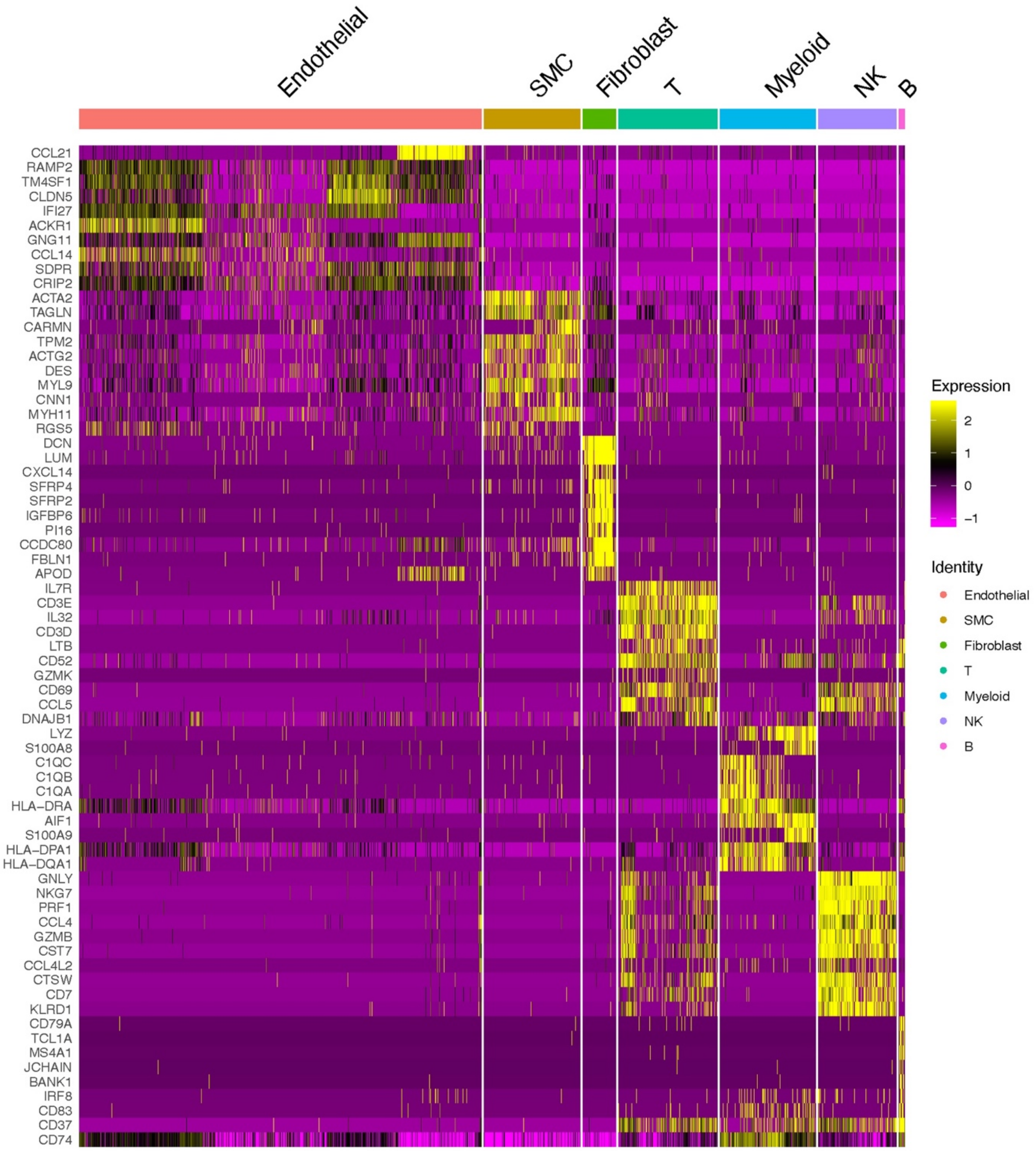
Endothelial
SMC
Fibroblast
T
Myeloid
NK
B
CCL21
RAMP2
TM4SF1
CLDN5
IFI27
ACKR1
GNG11
CCL14
SDPR
CRIP2
ACTA2
TAGLN
CARMN
TPM2
ACTG2
DES
MYL9
CNN1
MYH11
RGS5
DCN
LUM
CXCL14
SFRP4
SFRP2
IGFBP6
PI16
CCDC80
FBLN1
APOD
IL7R
CD3E
IL32
CD3D
LTB
CD52
GZMK
CD69
CCL5
DNAJB1
LYZ
S100A8
C1QC
C1QB
C1QA
HLA−DRA
AIF1
S100A9
HLA−DPA1
HLA−DQA1
GNLY
NKG7
PRF1
CCL4
GZMB
CST7
CCL4L2
CTSW
CD7
KLRD1
CD79A
TCL1A
MS4A1
JCHAIN
BANK1
IRF8
CD83
CD37
CD74
Expression
2
1
0
−1
Identity
Endothelial
SMC
Fibroblast
T
Myeloid
NK
B

**S Fig. 3: Cell clusters in leiomyomas** A) UMAP showing different cell clusters present in uterine leiomyomas B) Heatmap showing the top 10 genes used for cluster identification. Columns denote cells, rows denote genes.

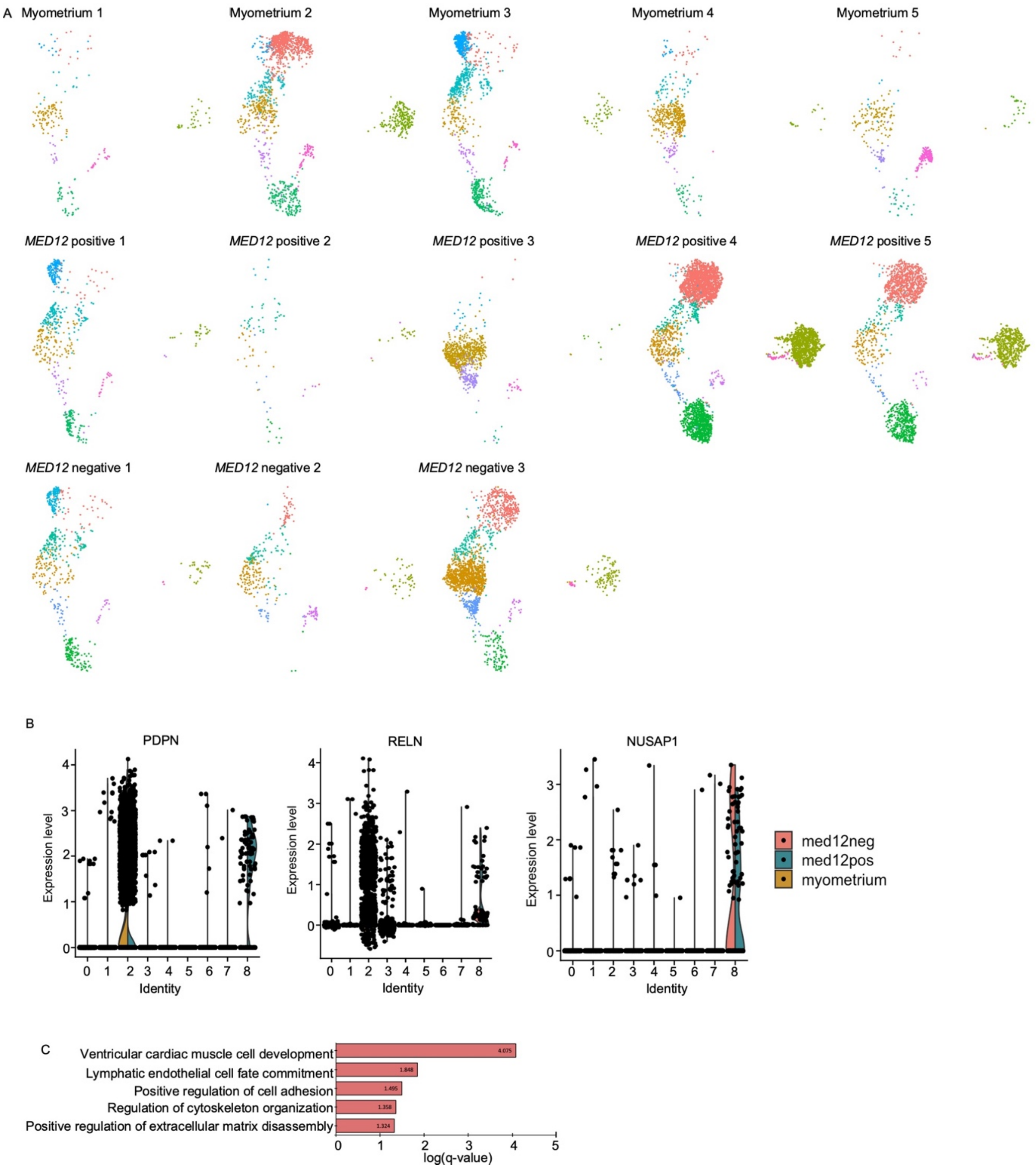
A
Myometrium 1
Myometrium 2
Myometrium 3
Myometrium 4
Myometrium 5
MED12 positive 1
MED12 positive 2
MED12 positive 3
MED12 positive 4
MED12 positive 5
MED12 negative 1
MED12 negative 2
MED12 negative 3
B
PDPN
RELN
NUSAP1
Expression level
Identity
0 1 2 3 4 5 6 7 8
med12neg
med12pos
myometrium
C
Ventricular cardiac muscle cell development
Lymphatic endothelial cell fate commitment
Positive regulation of cell adhesion
Regulation of cytoskeleton organization
Positive regulation of extracellular matrix disassembly
4.075
1.848
1.495
1.358
1.324
0 1 2 3 4 5
log(q-value)

**S Fig. 4: Lineage identification of lymphatic endothelial cells in leiomyomas** A) UMAP showing the endothelial cell cluster per patient sample. B) Violin plot showing the expression of *PDPN, RELN* and *NUSAP1* in endothelial cell clusters of all three different sample conditions. C) GO analysis of DE genes showing enrichment in the expanded clusters in *MED12* variant positive leiomyomas and *MED12* variant negative leiomyomas compared to the normal myometrium respectively.

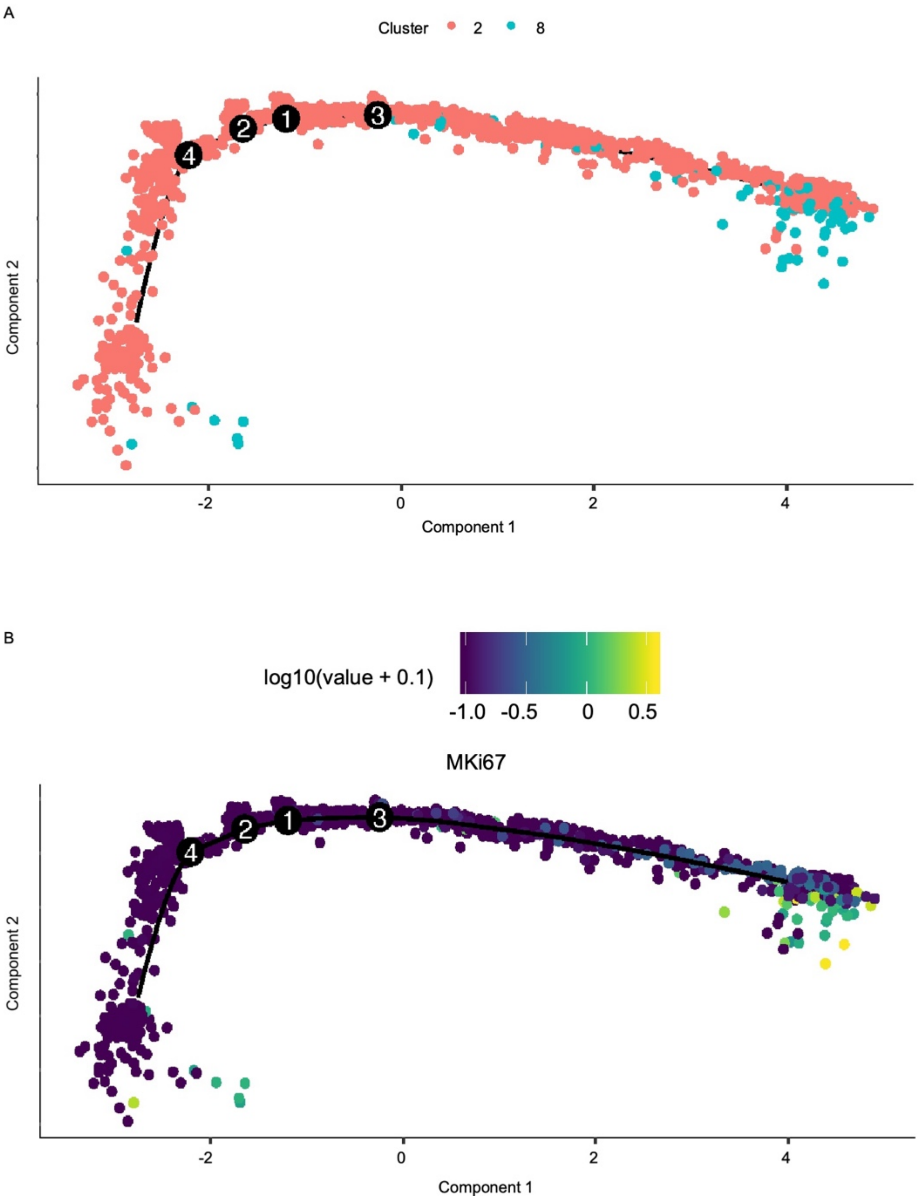


**S Fig. 5: Single-cell trajectory analysis of lymphatic endothelial cells using Monocle 2.** A) Pseudotime analysis of lymphatic endothelial cells. The cells on the tree are colored by clusters. B) log10 expression of MKi67 on the trajectory plot.

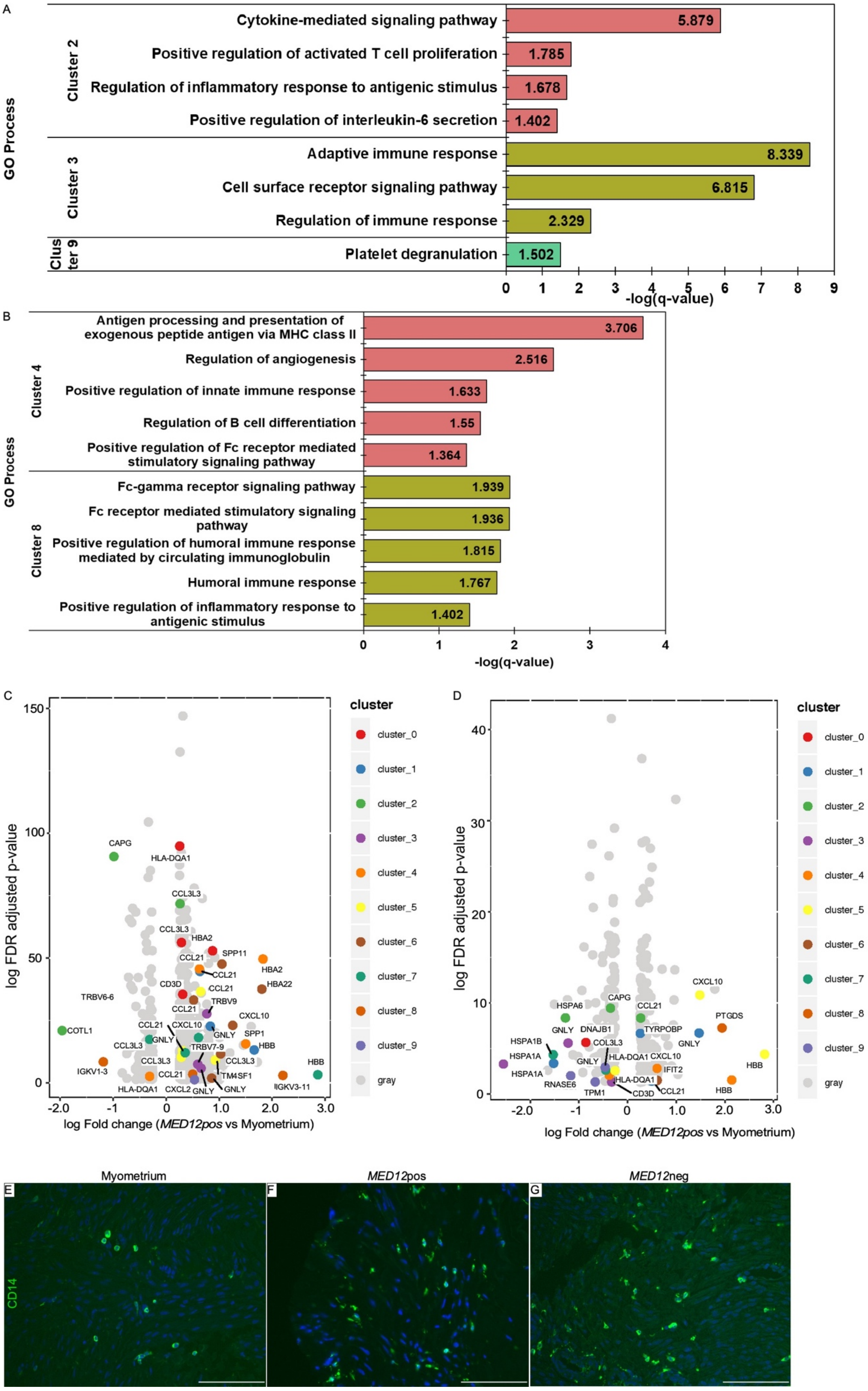

A
GO Process
Cluster 2
Cytokine-mediated signaling pathway 5.879
Positive regulation of activated T cell proliferation 1.785
Regulation of inflammatory response to antigenic stimulus 1.678
Positive regulation of interleukin-6 secretion 1.402
Cluster 3
Adaptive immune response 8.339
Cell surface receptor signaling pathway 6.815
Regulation of immune response 2.329
Clus ter 9
Platelet degranulation 1.502
0 1 2 3 4 5 6 7 8 9
-log(q-value)
B
GO Process
Cluster 4
Antigen processing and presentation of exogenous peptide antigen via MHC class II 3.706
Regulation of angiogenesis 2.516
Positive regulation of innate immune response 1.633
Regulation of B cell differentiation 1.55
Positive regulation of Fc receptor mediated stimulatory signaling pathway 1.364
Cluster 8
Fc-gamma receptor signaling pathway 1.939
Fc receptor mediated stimulatory signaling pathway 1.936
Positive regulation of humoral immune response mediated by circulating immunoglobulin 1.815
Humoral immune response 1.767
Positive regulation of inflammatory response to antigenic stimulus 1.402
0 1 2 3 4
-log(q-value)
C
log FDR adjusted p-value
log Fold change (MED12pos vs Myometrium)
cluster
cluster_0
cluster_1
cluster_2
cluster_3
cluster_4
cluster_5
cluster_6
cluster_7
cluster_8
cluster_9
gray
D
log FDR adjusted p-value
log Fold change (MED12pos vs Myometrium)
cluster
Myometrium
MED12pos
MED12neg
E
F
G
CD14

**S Fig. 6: Transcriptional changes in immune cell clusters in leiomyomas compared to the normal myometrium** A) GO enrichment analysis of the immune cells present in *MED12* variant positive leiomyomas compared to the normal myometrium B) GO analysis of the immune cell clusters expanded in *MED12* variant negative leiomyomas compared to the myometrium C-D) Volcano plots showing the log fold change in top genes in *MED12* variant positive leiomyomas compared to the myometrium and *MED12* variant negative leiomyomas compared to myometrium respectively. E) CD14 immunostaining in myometrium, *MED12* variant positive leiomyomas, *MED12* variant negative leiomyomas showing increased presence of macrophages in *MED12* variant negative leiomyomas.
